# Species demography, not metabolic rate, predicts community dominance independently of initial evenness

**DOI:** 10.64898/2026.09.24.754014

**Authors:** Daniela Almeida, Giulia Ghedini

**Author notes:** **Email(s) for correspondence:**, **Address for correspondence:** Daniela Almeida, GIMM Rua Quinta Grande 6, 2780-156 Oeiras, Portugal.

## Abstract

Understanding how species traits shape community properties is a central goal in ecology. But whether traits drive consistent patterns of dominance and assembly regardless of initial species abundances (evenness) remains unclear. We addressed this question using phytoplankton microcosm experiments to disentangle the effects of evenness and traits on community assembly and functioning, focusing on traits related to energy use (photosynthesis, size) alongside demographic parameters. Community assembly was highly deterministic, indicating that initial evenness had little influence on species dominance and final community structure. Instead, evenness drove community functioning. Demography was a stronger predictor of dominance than photosynthetic rates or individual size, suggesting that growth and competitive success depend on multiple metabolic processes that cannot be reduced to simpler proxies. These results show that community structure can emerge predictably across a range of initial conditions, and highlight the importance of demographic metrics over simpler metabolic descriptors for forecasting community outcomes.

## Introduction

Is community structure predictable? This question has long interested ecologists, but predictability remains debated. Community structure should be predictable from how species perform in isolation if species traits predominantly determine species performance and rankings (Arim *et al*. 2023; Vellend *et al*. 2014). Experiments on microbes provide some support for this idea, showing that community assembly is often predictable from the bottom-up by studying how species perform alone and in pairs, at least when diversity is low (Fant *et al*. 2026; Friedman *et al*. 2017a; Lee *et al*. 2023; Vandermeer 1969). This predictability suggests that traits linked to growth (e.g. size or metabolic rates) might be strong determinants of a species’ success in a community. A global analysis on plants, for instance, shows that traits such as wood density, specific leaf area and maximum height, consistently influence competitive interactions (Kunstler *et al*. 2016). But studies on other primary producers, such as phytoplankton, have not shown clear correlations between traits (e.g., cell size) and competitive outcomes (Fant *et al*. 2026; Gallego *et al*. 2019). More generally, which traits consistently inform on community assembly is an open question (Herben & Goldberg 2014; Levine *et al*. 2025) because their effects can be influenced by species interactions and environmental conditions (Adler *et al*. 2013; Chauvet *et al*. 2017; Laughlin *et al*. 2020).

In particular, the effects of traits on species performance can be mediated by the density and identity of neighbours (Buche *et al*. 2026). In other words, trait effects depend on the community in which a species is embedded (Su & Callaway 2026). This context-dependency raises an important challenge given that variation in species densities is a common feature of communities, reflected by differences in evenness (or the complementary term, dominance) (Hillebrand *et al*. 2008).

The effects of evenness on communities have mostly been studied from the perspective of community functioning. Evenness reflects the identity of dominant traits and their distribution in a community (Hillebrand *et al*. 2008). When species have similar abundances, complementarity effects can be enhanced leading to a positive relationship between functioning and evenness (Zhou *et al*. 2026). Through these same effects (i.e. by changing species interactions), evenness can also alter community assembly but this aspect has received less attention (Ehsani *et al*. 2018). Studies of community assembly often focus on coexistence, i.e. whether species are present or absent, without placing emphasis on their relative abundances – either in the initial or final community (Friedman *et al*. 2017b; Meroz *et al*. 2021). Yet, initial differences in species’ densities can affect the performance of species in a given community, particularly when species display trade-offs between intrinsic growth rates and the ability to tolerate competition (Buche *et al*. 2026; Martorell & Freckleton 2014).

We use microcosm experiments on marine phytoplankton to disentangle the importance of species traits and evenness on community assembly and functioning. Phytoplankton are photosynthetic microbes that mimic the tractability of bacterial systems, whilst enabling tests of trait-based approaches given their size diversity (Finkel *et al*. 2010). Metabolic rates (respiration, photosynthesis) and nutrient affinity scale allometrically with phytoplankton cell size while growth rates follow a unimodal relationship (Hillebrand *et al*. 2022; Wickman *et al*. 2024). Therefore, we use species that vary in cell volume by about two orders of magnitude to explore a range of species performance and competitive ability. In this system, community assembly can be quantitatively predicted (i.e. final species abundances) based on the demographic parameters of species in isolation and their interactions in pairs – across communities of different geographic origin and exposed to different environmental conditions (Fant *et al*. 2026). But whether variation in initial species densities affects community assembly is untested. Given that the density and identity of competitors can mediate the strength and direction of species interactions (Buche *et al*. 2026; Hillebrand *et al*. 2008), communities starting with different evenness might reach different endpoints and might function at different rates.

To answer this question, we assembled communities composed by the same pool of five species, and varied their initial abundance and the identity of dominant species. We then ask 1) how variation in initial evenness influences the predictability of community assembly, 2) which species traits predict dominance, and 3) whether the traits of the dominant species or the balance of species have stronger effects on functioning (Figure 1). To do so, we tracked the trajectory of communities over time quantifying changes in their structure (species abundances) and functioning (community biomass production and metabolic rates). To determine the role of species traits on community dynamics, we measured species characteristics at two levels: the demographic metrics of populations that capture the net effects of traits on performance (Laughlin *et al*. 2020), and two organismal traits related to energy use and growth: a physiological rate (photosynthesis) and a morphological trait (individual size) (Fant & Ghedini 2024; Levine *et al*. 2025). Our analysis of demographic metrics was done in two steps. First, we quantified the intrinsic growth rate (*r*) and maximum biovolume (K) of each species in monoculture and their performance in communities. Then we grew each monoculture under a resource gradient to identify demographic trade-offs in their ability to grow (*r*), tolerate competition (intraspecific competition coefficient, *α*_ii_), and the combined effects of these parameters on carrying capacity (K) (Marshall *et al*. 2023). Exploring demographic trade-offs allows us to clarify dominance patterns in communities where species compete for limited resources (Buche *et al*. 2026).

**Figure 1.**
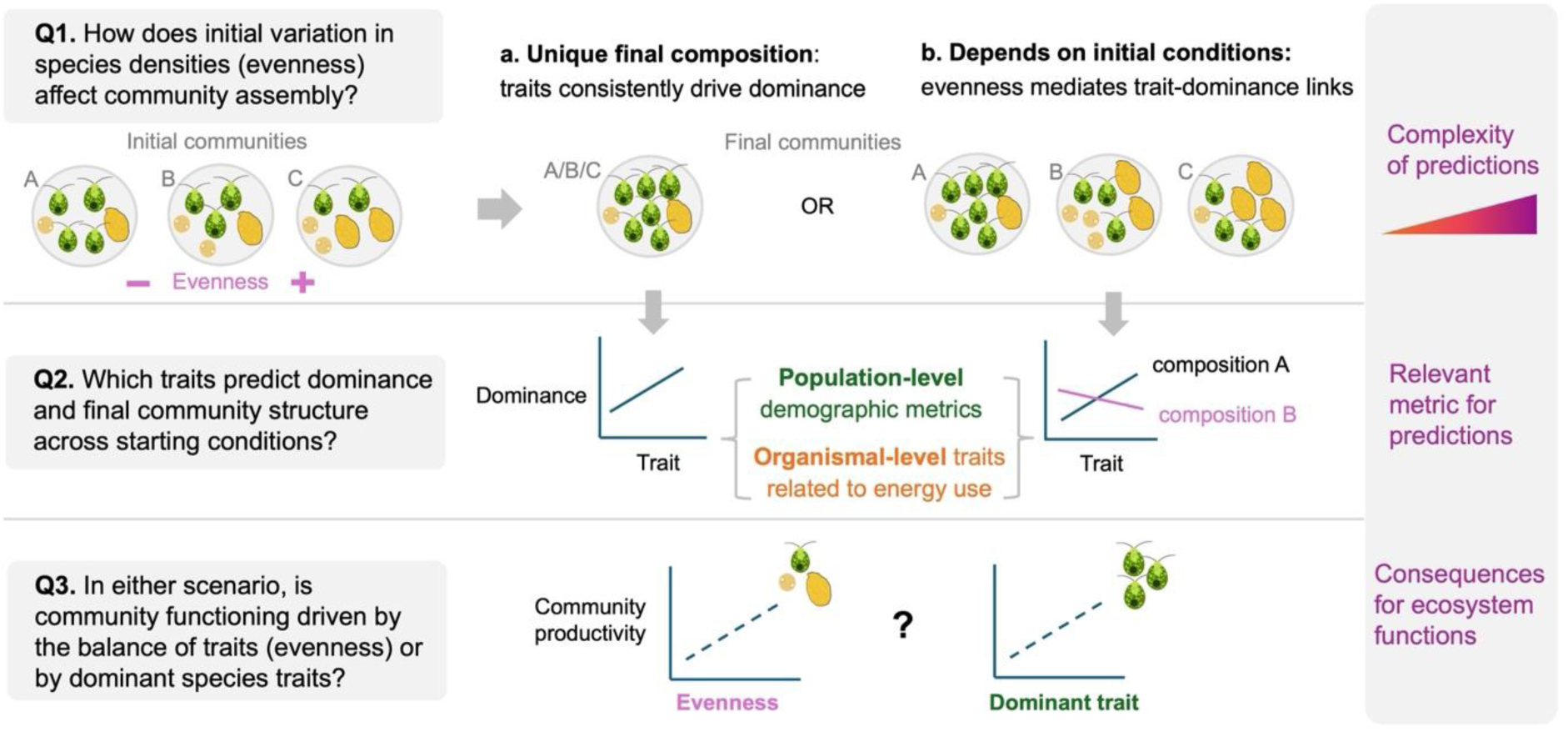
Conceptual overview. We use phytoplankton communities of varying evenness (including differences in the identity of dominant species) to determine Q1) how evenness affects the predictability of community assembly. If trait effects dominate, differences in species performance should lead to a consistent final composition, independently of the initial conditions (a). If variation in species densities (evenness) affects performance, final composition depends on the initial conditions (b). For either scenario we ask Q2) which of species traits best predict community outcomes, measuring traits at two levels: demographic parameters that capture the effects of traits on fitness and organismal traits related to energy use (photosynthesis rate, cell size). Finally, we investigate the functional consequences of dominance, specifically asking Q3) if community functioning is more strongly driven by the balance of species traits or by the identity of dominant traits.

Our hypotheses were that: H1) communities of different evenness would assemble to different final structures (i.e., species abundances measured as biovolume, a proxy for biomass) because variation in species densities affects performance; H2) organismal traits and demographic parameters related to growth (size, photosynthesis rate, intrinsic growth rates) would be good predictors of species’ success in a community but their predictive power would decrease in uneven communities where final composition might be more strongly influenced by initial species dominance; H3) community productivity would be more strongly driven by evenness (“complementarity effects”) than by the identity of dominant species (“selection effects”).

## Methods

### Testing how evenness affects community assembly

We established six community compositions based on the relative abundance of species and the identity of the dominant species, replicating each of them five times. The evenness of these communities, measured as Pielou’s Evenness Index (J), was chosen a priori to be one of four levels, and the two intermediate levels of evenness were repeated with two compositions (Table S1). All communities were composed of five species sourced from the Roscoff Culture Collection (France): *Dunaliella tertiolecta*, *Tisochrysis lutea*, *Nannochloropsis granulata*, *Phaeodactylum tricornutum* and *Amphidinium carterae*, encompassing a range of functional groups and traits (Table S2). Monocultures of each species were cultured in parallel (*n* = 5) to quantify demography and traits in isolation (described below). We established all cultures with an initial biovolume of 10^4^ μm^3^/μl. Samples were grown in tissue culture flasks (TC flask T75 standard, Sarstedt, Germany) set to 100 mL total volume, using artificial seawater F media (Guillard & Ryther 1962), kept in a temperature-controlled room at 18±1°C and under a 14-10 hr day:night cycle, with a light intensity of 70 μmol photons m^-2^ s^-1^. To understand community dynamics, we tracked the biovolume growth of the two dominant species identified in the communities (*Dunaliella* and *Nannochloropsis*) in a separate pairwise experiment (Supplementary Methods).

### Measuring cell size and abundance

We sampled all communities and monocultures over the course of 27 days (day 0, 4, 9, 14, 20, 27), until community biovolume was strongly dominated by one species (Figure S1). Each sampling day, we collected 10 mL from each culture and replaced it with 10 mL fresh media. The 10 mL sample was used to determine species densities, size, and biovolume using microscopy and measure metabolic rates.

We fixed a 1000 μl sample with 10% lugol and loaded 10 μl onto a cell counter (Neubauer improved chamber, MARIENFELD) to take 20 photos at 400× using an inverted microscope (Evident Scientific IX73). For the first and last sampling days we quadrupled the effort for a more accurate estimate of species abundances. We used Fiji (Schindelin *et al*. 2012), an open-source software for image analysis, to measure the number of cells (cells/μl), their length and width which were used to calculate the average cell volume by assigning an approximate geometric shape to each species (Table S2). Biovolume (μm^3^/μl) was calculated as the product of cell number and size.

### Measuring metabolic rates

We monitored changes in oxygen levels of monocultures and communities under the light (∼25 minutes) and dark (∼45 minutes) to measure photosynthesis and dark respiration rates, respectively. We measured more frequently in the early growth stages and less frequently as growth started to slow down: every day in week 1, every other day in weeks 2-3, twice in week 4, and once on the last experiment day for a total of 16 sampling points. Given the frequency of metabolic measurements, we used optical density as a measure of total biovolume (instead of microscopy which is substantially more time-demanding). We measured absorbance at 450 nm wavelength (Multiskan sky, firmware version 1.0.58, UI version 1.51.32.0) and corrected it with blanks made of the artificial seawater F media.

Metabolic rates were measured on 5 mL samples placed in glass vials, loaded onto 24-channel readers (SDR SensorDish® Reader, software version SDR_v4.0.0) which were previously calibrated at 0% and 100% oxygen levels. In each vial, we added 50 µl of sodium bicarbonate solution (1% of the vial volume) set to a 2 mM concentration to avoid carbon limitation. Because our cultures were not axenic, we established three blanks per treatment by centrifuging the algal cells in the samples down (5000 rpm for 10 minutes) and using the obtained supernatant to correct for bacterial respiration. The photosynthesis and respiration rates (μmol O_2_/min) of each sample were calculated as:

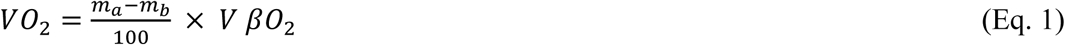

where m_a_ is the rate of change in oxygen levels of the sample (min^-1^), m_b_ is the mean O_2_ level of the respective blanks (min^-1^), V is the sample volume (0.005 L) and βO_2_ is the O_2_ capacity of air-saturated seawater at ∼20℃ and 35 ppt salinity (225 µmol O_2_/L). The first five minutes of oxygen readings during the light phase were discarded to account for acclimation to the light conditions. Similarly, we discarded the first 20 minutes of oxygen consumption in the dark to allow species to fully acclimate to the dark conditions.

### Assessing demographic trade-offs across resource levels

To quantify trade-offs in species performance, we grew each monoculture on a gradient of nutrient concentrations corresponding to 100%, 60%, 30% and 10% of the artificial sewater F media used above (*n* = 3 per combination of species and nutrient concentration). We grew species in 100 mL cell culture, except for *Amphidinium* for which we used 60 mL (because its growth was very slow and we did not have enough culture to make 100 mL). Initial biovolume was standardised at 4 × 10^4^ μm^3^/μl and species were grown under the same temperature and light conditions described above. We tracked the number, size, and biovolume of each species over time using microscopy, as described earlier, until carrying capacity for a maximum of 37 days (1, 2, 3, 7, 9, 10, 14, 16, 18, 23, 25, 28, 32, 37). We used a Lotka-Volterra growth model with explicit intraspecific competition coefficient (α_ii_) to explore the changes in intrinsic growth rates and density-dependence, and their combined effects on carrying capacity (Fronhofer *et al*. 2023; Mallet 2012; Marshall *et al*. 2023). Specifically, we fit the model below to the time series of biovolume data (μm^3^/μl) of each replicate:

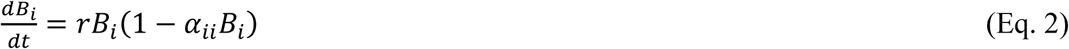

where *B_i_* is the biovolume of species *i*, *r* is the intrinsic rate of increase, and *α_ii_* is the intraspecific competition coefficient (which is also 1/K, the carrying capacity).

### Statistical analyses

All statistical analyses were performed in R studio, version 2025.09.2+418.

#### H1. Testing how evenness influences community assembly

*Changes in community structure*: We created a Bray-Curtis dissimilarity matrix on relative species biovolume data for the two timepoints of interest (initial = day 0, final = day 27). We then tested differences in community structure, including sampling time (initial vs final) and community composition (6 levels, Table S1) as categorical predictors using a permanova model (adonis2) from the *vegan* package. We tested for significant effects of sampling time, initial evenness, and community composition using the envfit() function and used a PCoA plot to visualize similarities across communities.

*Changes in evenness*: We calculated the change in species evenness (Pielou’s J) for each community as the difference between the final (day 27) and initial evenness (day 0), based on biovolume. For day 0, we used the “observed evenness” (calculated from measured species biovolumes) rather than the “expected evenness” decided a priori (Table S1) to account for any experimental error. We then used a linear model to test if evenness change was related to the initial evenness and community composition. In the model, we used expected evenness so that the predictor was not used for calculating evenness change.

*Species growth in communities*: Given that some species displayed a long lag phase or grew little in communities, we could not fit the r-*α* model (Eq. 2) to all species and replicates. Therefore, we used the model *fit_easylinear* from the R *growthrates* package to estimate species intrinsic growth rate (*r*) in the communities. The method uses a heuristic approach to determine maximum growth rates from the log-linear part of a growth curve, fitting linear regressions to all subset consecutive datapoints, from a defined number of points (h; we set h = 3 for all species) (Petzoldt 2022).

#### H2. Determine which species characteristics predict dominance

To explore which species characteristics predict dominance in communities, we tested how demographic parameters (*r_mono_*, K*_mono_*) and morphological (cell size) or physiological traits (photosynthesis rate per unit biovolume) measured in monoculture relate to the final abundance of species in communities (in %, i.e. species dominance) using a linear model for each predictor. Dominance, cell size, and photosynthesis were log_10_-transformed to improve normality and all models included an interaction between predictors and initial evenness.

The morphological trait cell size was determined as the average cell volume of each monoculture replicate during the entire growth curve. The demographic parameter K*_mono_* was calculated as the maximum biovolume value reached during the experiment, for each replicate. For the intrinsic rate of biovolume increase (*r_mono_*) we used the approach described above for communities (*fit_easylinear* model; h = 4). For two monoculture samples of *Phaeodactylum* and one for *Tisochrysis*, the model did not work, so we calculated *r_mono_* as:

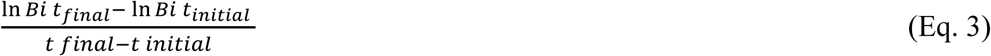

where B_i_ t_final_ and B_i_ t_initial_, correspond to the biovolume of the species at the end (day 27) and the beginning (day 9) of the exponential phase, respectively.

Photosynthesis rates per unit biovolume were calculated at two biovolumes that overlapped for all species (10^4.7^ and 10^5^ μm^3^/μl). The rescaling to a common biovolume was done because photosynthesis is sensitive to biomass density and species biomass (measured as either optical density or biovolume) did not fully overlap between species, which could lead to inaccurate comparisons (Marshall 2024). The rescaling uses the relationship between monoculture photosynthesis rates and biovolume to calculate the expected population photosynthesis rate at a fixed biovolume density in the centre of the range. Then we calculated rates *per cell* (diving the population rate by the total number of cells in the sample) and per unit biovolume (dividing *per cell* rates by the average monoculture cell size, averaged across all replicates). See Supplementary Information for more details.

*Species demographic trade-offs across resource levels*: To test how growth rates and sensitivity to competition varied with resource availability, we used linear models with *r* or *α*_ii_ calculated using the Lotka-Volterra model (Eq. 2) as response variable and resource concentration and species ID as predictors. To quantify trade-offs in performance we tested how the relationship between K (K = 1/*α*_ii_) and *r* varied between species, using a linear model with K as response variable, *r* and its interaction with species ID as predictors. To ensure that there are no systematic differences caused by the approach used, we compared estimates of *r* from Eq. 2 to those obtained using the *fit_easylinear* model and found no significant difference (paired t-test: t = 0.16, df = 47, p-value = 0.87).

*Validating the importance of demography for community outcomes*: We implemented a simple growth model to calculate the final biovolume of each species in the community (at day 27, *B_ifinal_*) based on its initial biovolume in the community (*B_iinitial_*) and its monoculure intrinsic growth rate (*r_mono_*; assuming that each species grows exponentially with no interactions):

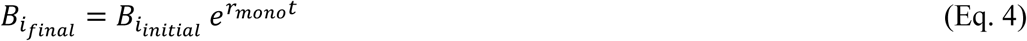

or including the monoculture carrying capacity (K*_mono_*, assuming only intraspecific self-limitation):

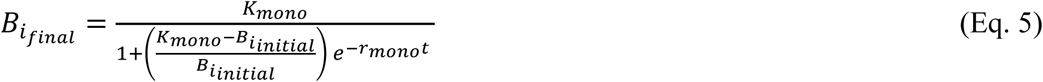

We plotted these predictions against observed final abundances, and calculated the residuals to explore where predictions deviated the most.

#### H3. Testing whether evenness or dominant species traits drive functioning

We tested whether initial evenness was a better indicator of community biovolume *r* (estimated using the *fit_easylinear* model) than the average monoculture growth rates weighted by their initial abundances (community weighted mean, CWM_r_mono_). To determine this, we compared the adjusted R^2^ of three linear models: including only initial evenness as predictor, only CWM_r_mono_, or both. Similarly, we tested whether final evenness was a better indicator of community K (the max. biovolume value reached by each replicate during the experiment) than the average monoculture carrying capacity weighted by the final abundances (CWM_K_mono_).

To clarify the effects of evenness on productivity, we tested how species’ intrinsic growth rates varied as a function of their initial proportion (pooling the estimates of *r* of each species obtained from monocultures and communities, with 1 corresponding to proportion in monoculture), including an interaction with species ID using a linear model. Finally, we tested the effects of initial evenness on community photosynthesis and respiration rates (which should affect biovolume production), including community optical density (OD) as a covariate in a linear model. We restricted OD values to < 0.2 to have overlapping ranges between all communities (only one community reached higher OD values). We removed negative rates of photosynthesis and respiration because they are biologically impossible – this happens when rates are very low and lower than blanks – leaving 384 observations for photosynthesis and 275 for respiration.

## Results

### 1) Community structure is highly predictable and mostly robust to variation in evenness

The structure of communities by the end of the experiment diverged significantly from initial conditions (Fig. 2) and converged in five out of the six compositions (Fig. 3a; sampling time × composition: F_11, 48_ = 70.45, p = 0.001). The primary axis (PCo1, 41% variance) reflects a compositional shift towards communities dominated by *Dunaliella* (Kendall’s rank correlation: z = 10.09, p-value < 0.0001, tau = 0.90). The second axis (PCo2, 14% variance) instead separates communities along a gradient of cell size (weighted average), with negative values corresponding to communities dominated by small species (*Tisochrysis*) and positive values corresponding to communities dominated by larger species (*Amphidium*) (z = 4.96, p-value < 0.0001, tau = 0.44). Both sampling time (initial vs final day) (R^2^ = 0.36, p = 0.001) and initial composition (R^2^ = 0.36, p = 0.001) explained changes in community structure, while initial evenness did not (R^2^ = 0.06, p = 0.18).

**Figure 2.**
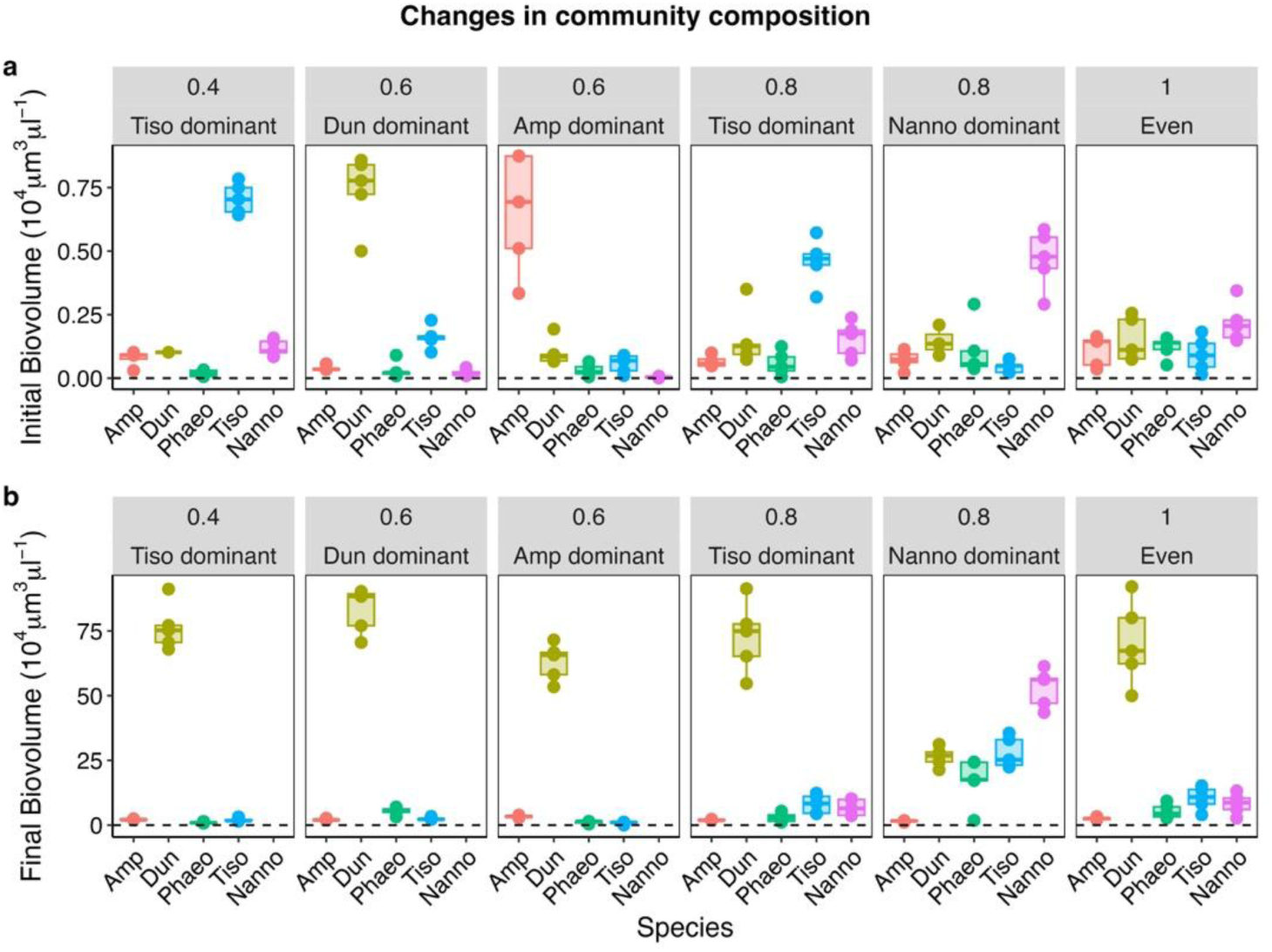
Absolute species abundances (biovolume) in each community at the start (a) and at the end of the experiment (b). Note different scale on y-axes. Species abbreviations: Amp = *Amphidinium*, Dun = *Dunaliella*, Phaeo = *Phaeodactylum*, Tiso = *Tisochrysis*, Nanno = *Nannochloropsis*.

**Figure 3.**
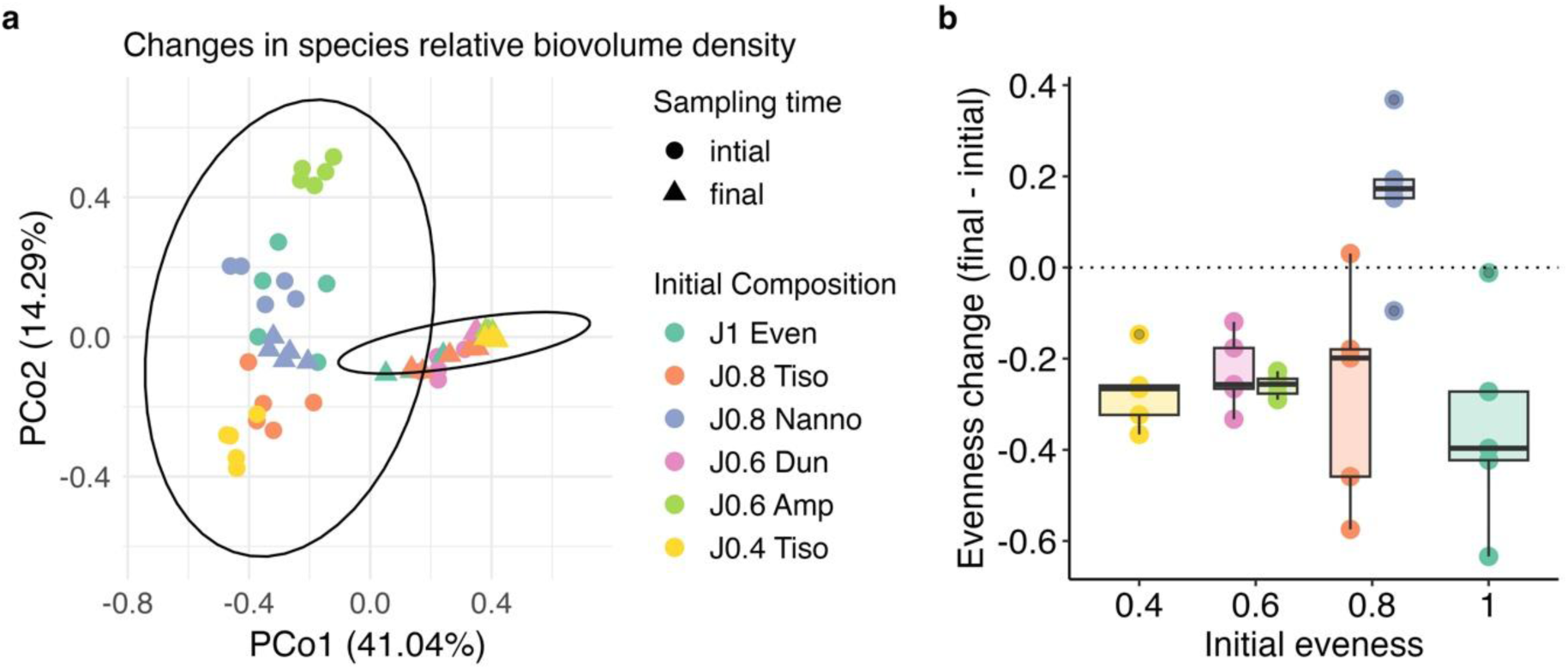
a) Principal coordinate analysis based on relative species biovolume shows that community structure changed significantly from the start to the end of the experiment (day 27), with nearly all communities converging to the same structure independently of initial conditions, except for the community initially dominated by *Nannochloropsis* (purple). b) This community was also the only one for which evenness increased over time – these changes in evenness were not explained by the initial evenness of communities.

Initial composition affected community assembly in two cases in particular: the community initially dominated by *Dunaliella* changed little over time and started with a composition that reflected the final composition of most communities (with *Dunaliella* strongly dominating biovolume and the other species present at low abundances) (Fig. 2, Fig. 3a). Instead, the community initially dominated by *Nannochloropsis* followed a different trajectory: it stayed close to its initial composition (*Nannochloropsis* dominated; Fig. 2) and increased in evenness over time (Fig. 3b). Changes in evenness were not explained by initial evenness but there was a significant effect of community composition (Fig. 3b; Table S3).

The pairwise dynamics between dominant species showed that *Dunaliella* was always the superior competitor of *Nannochloropsis* (Fig. S2). Therefore, the different trajectory of the community dominated by *Nannochloropsis* cannot be simply explained by its initial dominance. The higher abundance of subordinate species (Fig. 3b) seems a key factor to explain why *Dunaliella* did not dominate this community.

### 2) Demography explains dominance more than individual size or photosynthesis rates

Performance in monoculture varied substantially among species, including their intrinsic growth rates, duration of lag phase, and final biovolume (Fig. S3). The two dominant species in communities (*Dunaliella* and *Nannochloropsis*) were the most productive in monoculture, both in terms of intrinsic growth rates (*r_mono_*) and final biovolume (K*_mono_*). The intrinsic growth rate was the metric that most consistently and monotonically correlated with species dominance in communities (*r* effect: F_1,127_ = 142.58, p < 0.0001), while max. biovolume, cell size and photosynthesis per unit biovolume less so, all showing an interaction with evenness and non-monotonic relationships (Fig. 4, Table S4).

**Figure 4.**
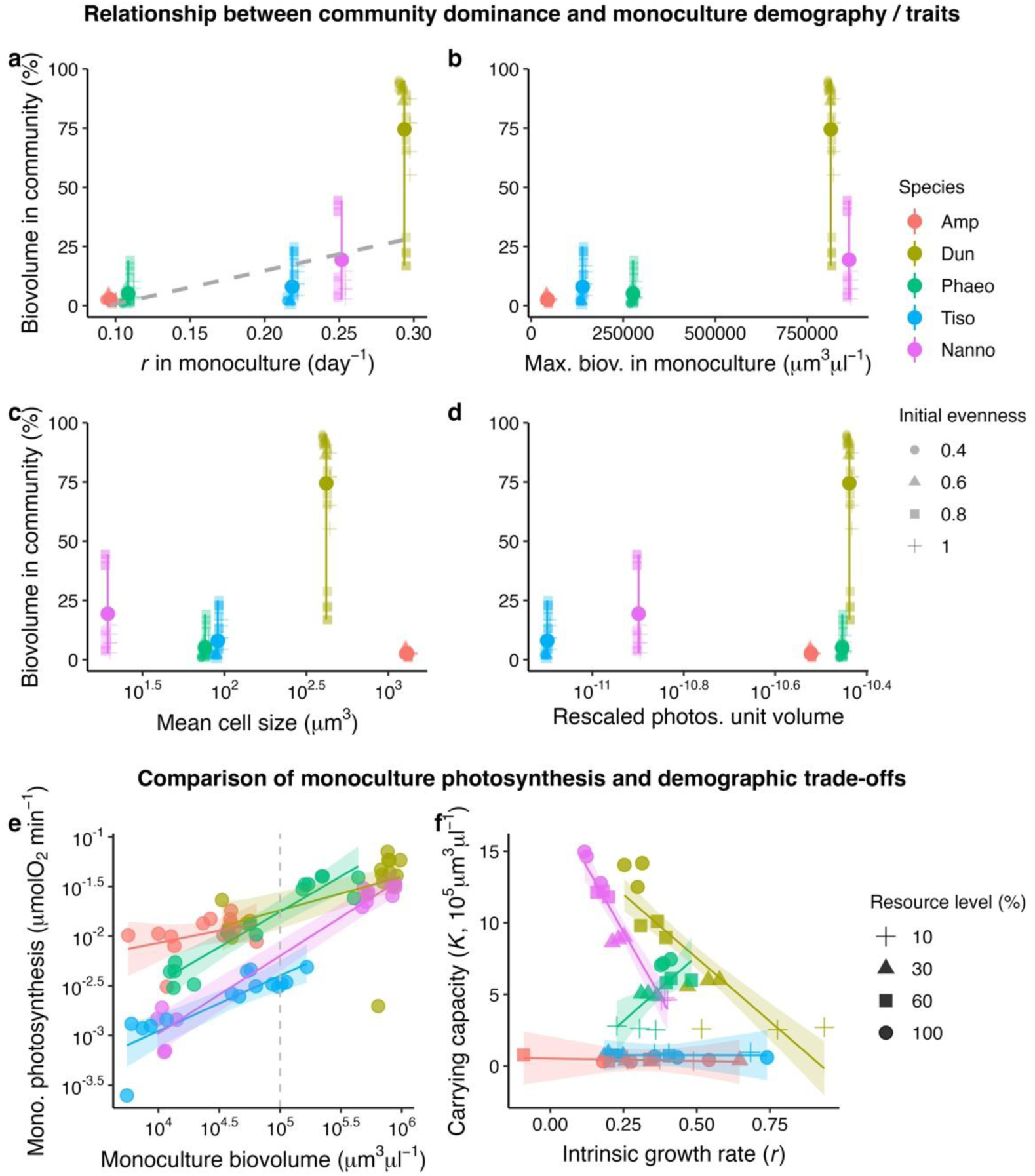
Species intrinsic growth rates in monoculture (a) were good predictors of dominance in the final communities, measured as % species biovolume (the grey line represents the predicted effect of *r* on dominance from the linear model). Instead, max. monoculture biovolume (b), cell size (c) and photosynthesis rates per unit biovolume (d) were less predictive. Photosynthesis rates per unit volume (d; μmol O_2_ min^-1^ μm^-3^) are calculated at a biovolume of 10^5^ μm^3^/μl for all species (grey vertical line in plot e). Predictions are not shown for b-d for clarity because all these predictors interacted with evenness and do not monotonically relate to dominance. Species differed much less in their photosynthesis rates (e) compared to their demographic traits (f), particularly when comparing photosynthesis at stadardised biovolumes (Figure S5). Assessing demographic parameters across resource levels showed that species had different growth responses and tolerance to competition as nutrient concentrations changed (Figure S7), leading to different trade-offs (or trade-ups) between *r* and K (f). The dominant species in communities (*Dunaliella*) was sitting at a Pareto optimality front between growth rates and carrying capacity (f).

Interspecific differences in photosynthesis rates were less obvious than differences in demography (Fig.4-f). Still, species varied in their metabolic density-dependence (i.e. the sensitivity of photosynthesis to intraspecific competition) as shown by differences in the slopes between monoculture photosynthesis and biomass (biovolume: Fig. 4g, Table S5; optical density: Fig. S4, Table S6). The dominant species (*Dunaliella*) did not have the steepest slopes (which would indicate less sensitivity to competition) but had higher intercepts. However, comparing species at a fixed biovolume better shows that differences in photosynthesis rates were subtle. At fixed biovolume (Fig. S5), per cell photosynthesis increased with cell size with a slope of 1.28, not significantly different from 1 (CI: 0.98, 1.57) as previously demonstrated (Fant & Ghedini 2024); a slope of ∼1 means that cells of each species produced energy at similar rates per unit biovolume (when compared at the same biomass density), explaining why photosynthesis rates are weak predictors of dominance in communities.

Conversely, species varied widely in their demographic performance and ability to grow under resource competition (Fig. 4f; Fig. S6). Dominant species (*Dunaliella* and *Nannochloropsis*) showed similar changes in *r* and *α*_ii_ as resource concentration decreased: their intrinsic growth rates increased at the cost of their sensitivity to competition (Fig. S7), reducing carrying capacity and leading to a clear *r*/K trade-off (Fig. 4f; Tables S7-S9). The three subordinate species showed different patterns: a positive relationship between *r* and K (*Phaeodactylum*), or a near flat relationship because changes in resource levels affected *r* but not so much the sensitivity to competition (*Tisochrysis*, *Amphidinium*). The dominant *Dunaliella* outperformed all species in its ability to grow at low resource levels and was sitting at Pareto optimality front of max. growth rate and carrying capacity (Fig. 4f).

To validate the importance of demographic parameters for community assembly, we used the intrinsic growth rate in monoculture (*r_mono_* in Fig. 4a) to predict the final species biovolume in each community. While this approach led to reasonable predictions considering its simplicity (Spearman’s correlation: p-value < 0.0001, ρ = 0.66), including carrying capacity (K*_mono_* in Fig. 4b) improved predictions across all starting conditions, particularly for the dominant species *Dunaliella* (p-value < 0.0001, ρ = 0.69; Fig. 5). This improvement shows that *Dunaliella* is little affected by interactions with other species, but the same does not hold for the other dominant species (*Nannochloropsis*) or the subordinate competitors (see large residuals in Fig. 5c particularly in more even communities).

**Figure 5.**
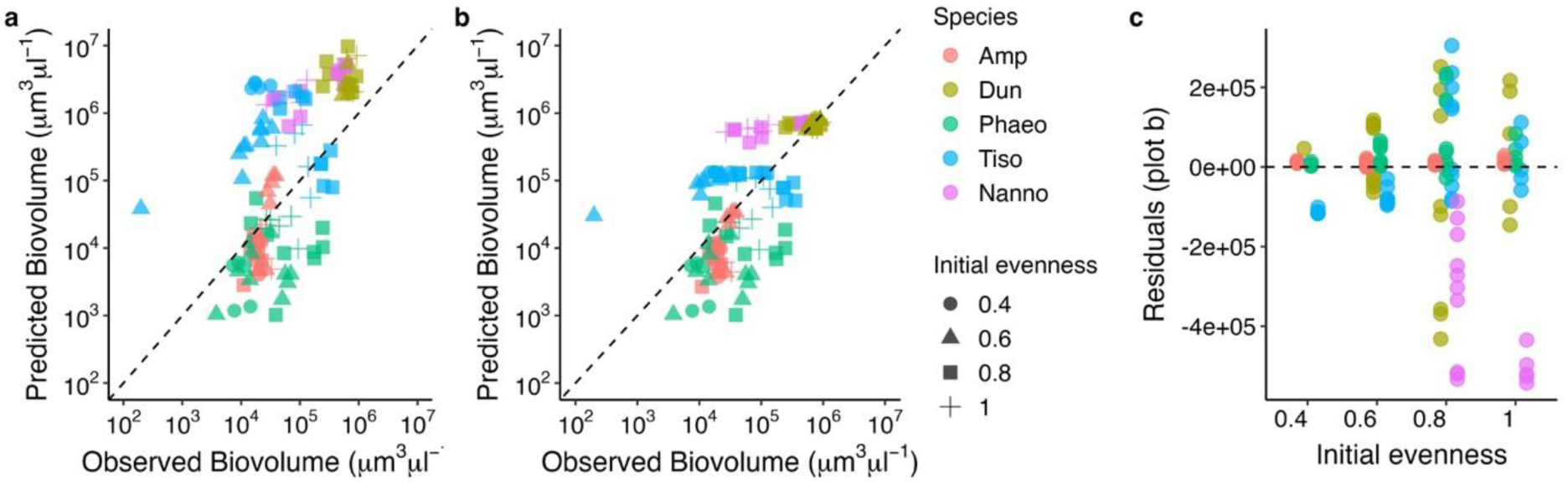
Final species abundances in communities can be predicted reasonably well from demographic parameters in monoculture across all starting conditions (evenness). Predictions based on monoculture intrinsic growth rates alone (a) are improved when accounting for carrying capacity (b), indicating that all species experience self-limitation in the community. The residuals of plot b show larger errors for more even communities (c), particularly for one of the dominant species, *Nannochloropsis*, indicating substantial differences in how the two dominant competitors (*Dunaliella*, *Nannochloropsis*) interact with other community members.

### 3) Evenness, more than dominant traits, drives community productivity

Community intrinsic growth rates (*r,* measured on biovolume) were positively related to both evenness (F_1, 28_ = 19.1, p = 0.0001) and dominant species characteristics (monoculture intrinsic growth rates weighted by their initial abundance; F_1, 28_ = 7.9, p = 0.009), but evenness was the stronger driver (Fig. 6a-b; adj. R^2^ evenness: 0.38; monoculture growth: 0.19, both: 0.55).

**Figure 6.**
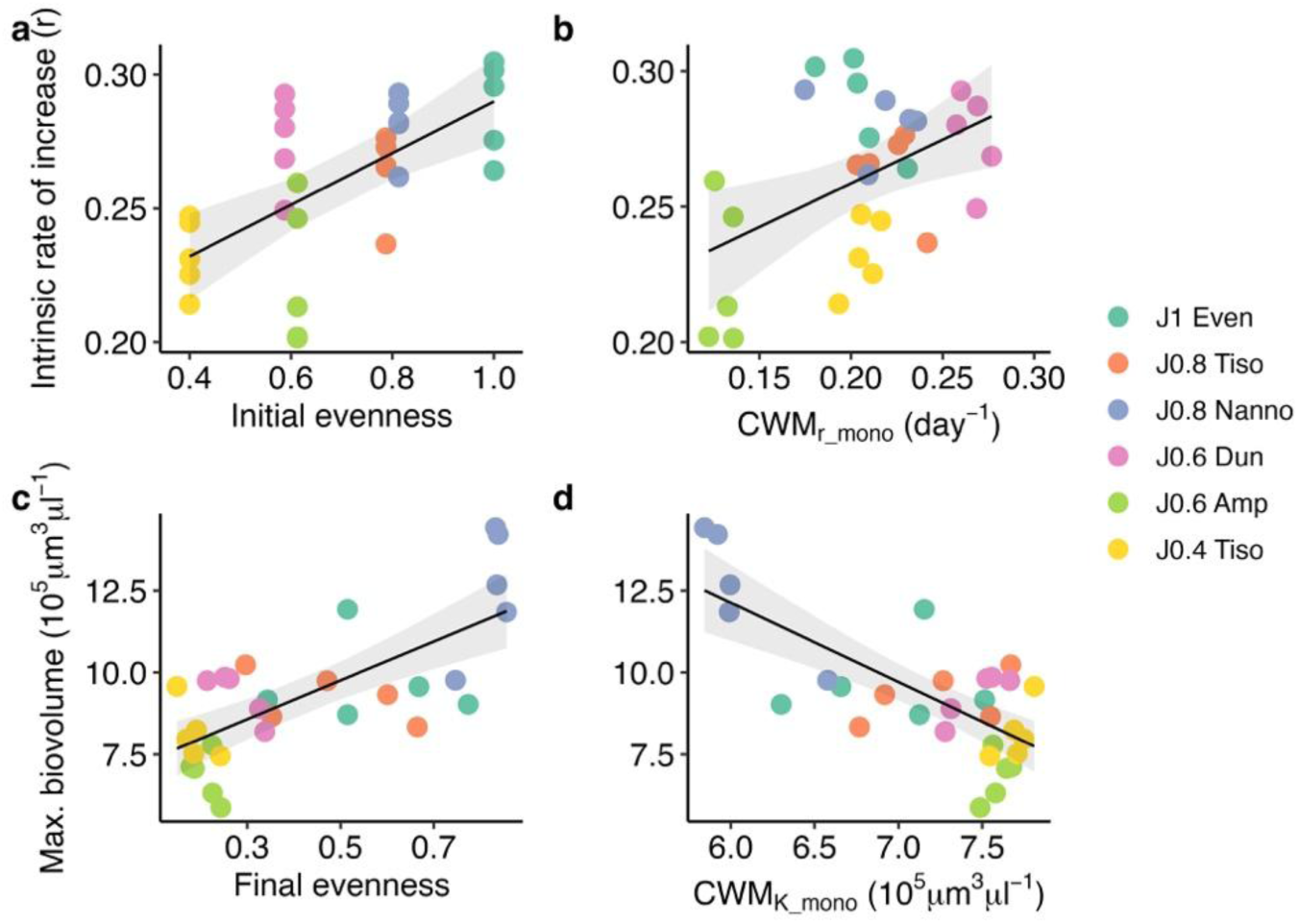
Communities’ intrinsic growth rate (biovolume) was positively related to both initial evenness (a) and the traits of dominant species (i.e. monoculture growth rates; b) but evenness was a stronger driver (adj R^2^ = 0.38 vs 0.19). Similarly, community maximum biovolume increased with final evenness, indicating complementarity effects (c), but declined in communities strongly dominated by species with high K in monoculture (d).

Max. community biovolume similarly increased with evenness (F_1, 28_ = 28.1, p < 0.0001), but was negatively related to monoculture K weighted by final species abundances (F_1, 28_ = 31.7, p < 0.0001) (Fig. 6c-d; adj. R^2^ evenness: 0.48, monoculture K: 0.51; both: 0.50). Note that we cannot exactly disentangle the effect of evenness from those of monoculture K because the two were strongly negatively correlated. Still, this shows that even communities, which had a lower weighted mean of monoculture carrying capacity, reached greater biovolumes than uneven communities strongly dominated by productive species (*Dunaliella*).

The positive effects of evenness on community productivity can be understood by assessing variation in species growth rates as a function of their initial proportion. Most species showed a negative correlation between *r* and their initial proportion, a sign that intraspecific competition is higher than interspecific competition, except for *Nannochloropsis* which showed the opposite pattern (Fig. S8, Table S10).

The effects of evenness on community biovolume were not visible on community photosynthesis rates (Fig. S9a; F_1, 367_ = 0.03, p = 0.86; Table S11). Community respiration instead increased more slowly with biomass (optical density) when evenness was high (F_1, 258_ = 5.87, p = 0.016; Table S11), but we have low confidence in this analysis given that these data are noisier (Fig. S9b).

## Discussion

Trait-based approaches to predict community assembly are lacking (Gallego & Narwani 2022; Zhang *et al*. 2024). In this study, we asked how initial differences in species abundance (evenness) affect the predictability of community outcomes in phytoplankton, given that species densities can affect the strength of intra- and inter-specific interactions (Hillebrand *et al*. 2008; Kardol *et al*. 2010). We further asked which species characteristics related to growth and energy use – namely demographic metrics, or physiological and morphological traits (photosynthesis rates, cell size) – can anticipate dominance patterns across different starting conditions and their consequences on functioning. We found that community assembly was highly predictable, (mostly) independently of initial evenness. Species abundances were not only extremely consistent between replicates but also converged in five of the six community compositions, despite initial differences in species proportions and dominance (contrary to H1).

These results suggest that deterministic processes driven by differences in species fitness (e.g., growth rates) had stronger impacts on community composition than any potential context-dependency driven by variation in species densities (e.g., advantage of rare *vs* disadvantage of abundant species, or changes in trait effects with competitor density) (Buche *et al*. 2026). While we observed convergence for most communities, one community type followed a completely different trajectory. This pattern seemed primarily driven by differences in how the two dominant species interacted with other community members more than with each other (as *Dunaliella* was always the stronger competitor in pairwise tests).

*Dunaliella* was little affected by interspecific interactions and strongly dominated communities reducing evenness (indeed we could predict its final biovolume remarkably well using only monoculture *r* and K); instead the abundance of *Nannochloropsis* was overestimated using this approach (Fig. 5), suggesting that this species was more constrained by interspecific interactions (as indicated by the positive relationship between its intrinsic growth rate and initial proportion, Fig. S8). We can speculate that the reduced growth of *Nannochloropsis* in the community allowed other community members to grow more evenly, preventing the dominance of a single species. Coexistence in multispecies communities can be therefore possible even when (for some species) intraspecific competition is weaker than interspecific competition, because community structure and higher-order feedbacks influence species trajectories and can override the simple two-species coexistence rule (Barabás *et al*. 2016).

Whether species traits consistently predict success in communities remains debated (Laughlin *et al*. 2020; Levine *et al*. 2017). Our results show that demographic metrics are better predictors of community dominance than organismal traits related to energy use (i.e. photosynthesis rates and cell size) across all starting conditions (only partially supporting H2), likely because demography captures the net effect of traits on fitness (Laughlin 2014; Wieczynski *et al*. 2021). These results are in line with studies on plants that show that “size or architectural traits” (such as plant height) have weaker effects on competitive differences than “growth or resource traits” (Ceballos-Núñez *et al*. 2025; Herben & Goldberg 2014).

Population growth is determined by many metabolic processes (Piorreck & Pohl 1984) which might not necessarily be captured by simpler descriptors of metabolism, such as photosynthesis rate or its relationship with size (Levine *et al*. 2025). Furthermore, demography can depart substantially from predictions based on metabolism-size relationships because metabolic rate (respiration or photosynthesis) often affects both resource supply (i.e. increasing competition) and acquisition (i.e. affecting growth) (Schuster *et al*. 2021). We can see some of these double effects of metabolism in our species. The two dominants (*Dunaliella*; *Nannochloropsis*) had the highest intrinsic growth rates in isolation and even increased them as nutrients declined. We can speculate that these species can increase max. growth rates under nutrient limitation via metabolic plasticity that allows a more efficient uptake or use of resources – these changes however come at a cost, reflected in higher competitive interactions (*α*_ii_) (Briddon *et al*. 2025; Marshall *et al*. 2023). Despite the resulting *r*/K trade-off, the dominant *Dunaliella* was sitting at a Pareto front that seemed to reflect the optimal compromise between intrinsic growth rate and competitive efficiency (Li *et al*. 2019), explaining its dominance in communities.

While species demography explained community assembly, functioning was more influenced by the diversity and balance of species traits (H3). By increasing trait diversity, evenness often facilitates complementarity effects (Daly *et al*. 2015; Zhang *et al*. 2012; Zhou *et al*. 2026). Indeed, most species grew faster when they occurred at lower initial proportions – a strong indicator of stabilising mechanisms that could aid both community productivity (Mariotte 2014) and coexistence (Desallais *et al*. 2026; Harpole & Suding 2007).

Our results might be dependent on the stable environmental conditions experienced by our cultures. Community structure might not be so deterministic under more variable environments: species traits are often plastic to environmental change and uneven species abundances might become more important under disturbance (Pérez-Ramos *et al*. 2019). Understanding how species traits, their plasticity, and proportions together modulate the predictability of community composition, and its effects on functioning, would be important to manage biodiversity in disturbed environments (Daly *et al*. 2015; Wang *et al*. 2023).

In summary, even though variation in species densities can affect performance and interactions (Buche *et al*. 2026), we find that deterministic differences in species fitness (demography) are the main drivers of community assembly, and this predictability is robust to variation in initial conditions (species abundances) – at least in the absence of disturbance. The assessment of demographic parameters revealed that strong competitors shared similar trade-offs between intrinsic growth rates and competitive efficiency. Simpler proxies for fitness, like photosynthesis rate and cell size, could not capture these demographic differences or explain variation in species dominance (see also (Fant *et al*. 2026; Gallego *et al*. 2019). Therefore, we should be cautious in extrapolating fitness from widely used morphological and physiological traits (Laughlin *et al*. 2020). Our results also reveal important limitations of demographic metrics: certain combinations of species led communities to suprisingly different compositions – with increased evenness and functioning. These effects could be understood by differences in how dominant species interacted within a community but were not captured by demographic parameters in monoculture. Therefore, despite the overall predictability of community assembly, an important component of diversity seems determined by more complex interactions, and their distribution, within communities (Barabás *et al*. 2016) that can make predictability more challenging.

## Acknowledgements

We are thankful to Ricardo Estevens for assistance during the experiments. We also thank Anna Lena Heinrichs and two anonymous reviewers for their feedback on the manuscript. DA was supported by an FCT Fellowship (UI/BD/154598/2023). This work was supported by an ERC Starting Grant by the European Union to G.G. (META_FUN, 101116029).

## Statement of authorship

DA and GG designed the study. DA performed the experiments and collected the data. DA and GG analysed the data and wrote the manuscript.

