## supplementary information for "Species demography, not metabolic rate, predicts community dominance independently of initial evenness"

**Supplementary Methods**

*Pairwise dynamics between the two dominant species*

To understand community trajectories, we tracked the biovolume growth of the two dominant species identified in the communities (*Dunaliella* and *Nannochloropsis*) in a separate pairwise experiment. We set up three mixtures of different initial biovolume densities (50:50, 80:20, 20:80) alongside the two monocultures in 30 mL cell culture flasks ($n$ = 2) keeping total biovolume constant ~$5 \times 10^4$ $\mu m^3/\mu l$. We tracked species biovolume using microscopy, as described in the main text, until growth stabilised (23 days).

*Rescaling of photosynthesis rates at a fixed biovolume density*

To compare species photosynthesis rates at the same biovolume density, we first obtained the relationship between photosynthesis and biovolume ($\mu m^3/\mu l$) for each species (both $log_{10}$-transformed), including an interaction between biovolume and species ID, using a linear model. Then we used this model to calculate the photosynthesis rate of each species at two fixed biovolumes in the centre of the range ($10^{4.7}$ and $10^5$ $\mu m^3/\mu l$). Finally, we calculated the photosynthesis rate per cell (dividing the rescaled population rates by the total number of cells in the sample) and estimated their relationship with cell size using a linear model ($log_{10}$-$log_{10}$ scale). To better visualise differences between species, we show photosynthesis rates per unit biovolume, calculated by dividing the rescaled per cell rates by the average size of each species (average of all replicates).

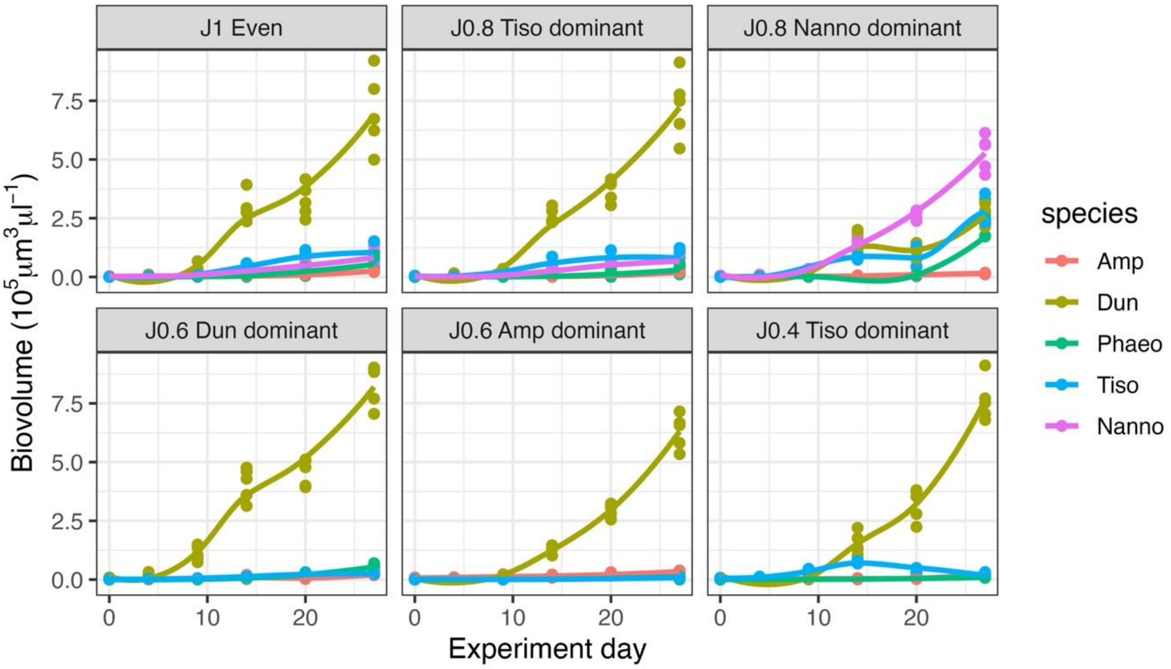


**Figure S1.** Species growth and dominance patterns were consistent in 5 out of the 6 community types, independently of initial evenness. Across all these communities, *Dunaliella* had the highest growth rates, dominating community biovolume within the first 10 days of the experiment. Only one of the six compositions (community dominated by *Nannochloropsis*) showed a different trajectory and final community structure. Facets legend: J indicates the initial evenness of the community (Pielou's J) followed by the identity of the dominant species. Species abbreviations: Amp = *Amphidinium*, Dun = *Dunaliella*, Phaeo = *Phaeodactylum*, Tiso = *Tisochrysis*, Nanno = *Nannochloropsis*.

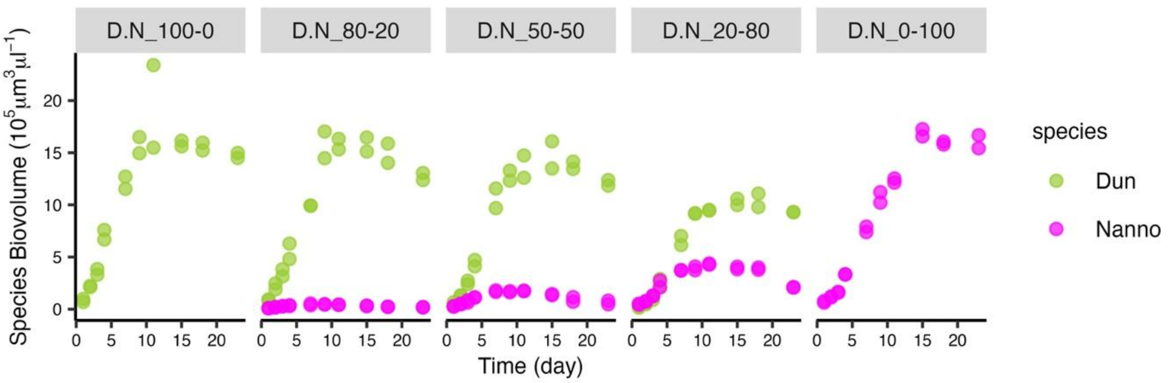


**Figure S2.** Pairwise dynamics between the two dominant species in communities, *Dunaliella* and *Nannochloropsis,* show that *Dunaliella* is always the superior competitor in pairwise competition, even when starting at a lower biovolume (20% *Dun* – 80% *Nanno*).

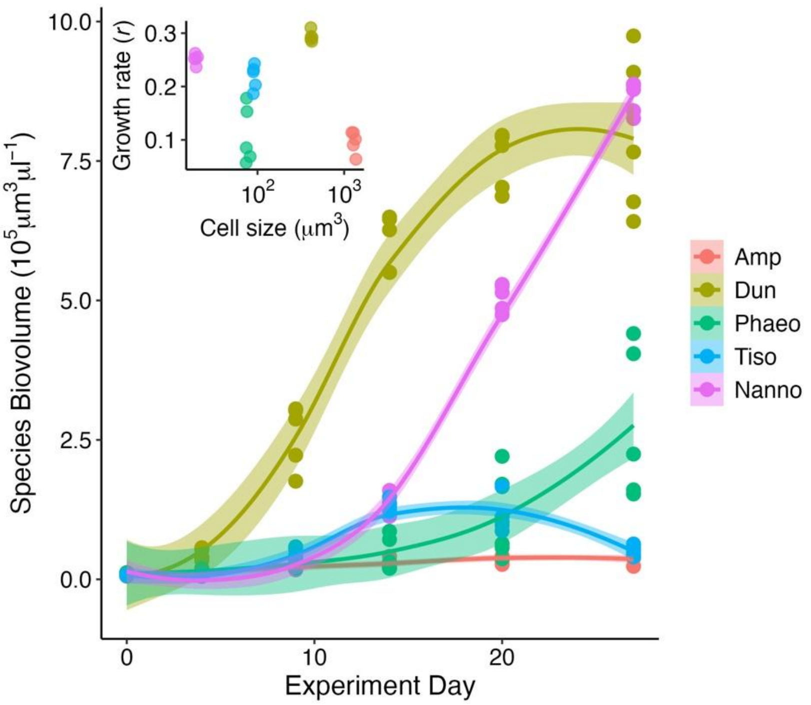


**Figure S3.** Species in monoculture showed variation in growth rates, lag phase, and max. biovolume. There was no clear relationship between intrinsic growth rate (*r*, calculated on biovolume) in monoculture and cell size (insert) but this should be explored over a larger size range.

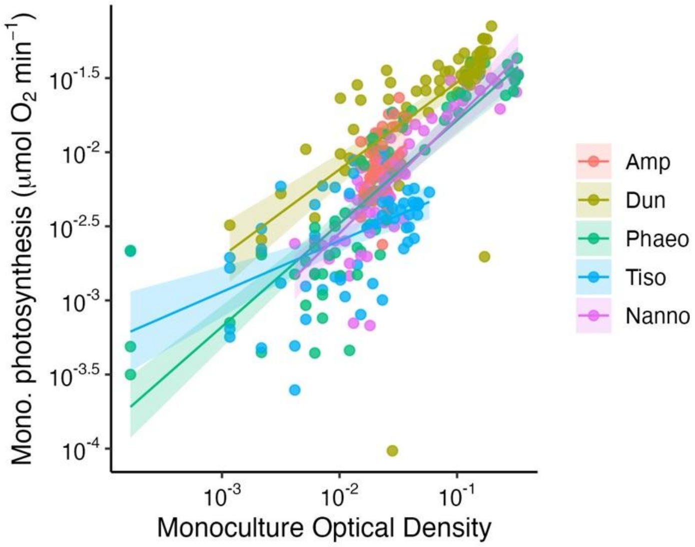


**Figure S4.** Monocultures showed some variation in photosynthesis rates, which increased with population optical density (OD) at different rates. The most dominant species, *Dunaliella*, also shows higher photosynthesis rates across the entire OD range. However, species optical density values were not fully overlapping – see Fig. S5 and 4 in main text for comparison of species photosynthesis rates at the same biovolume density.

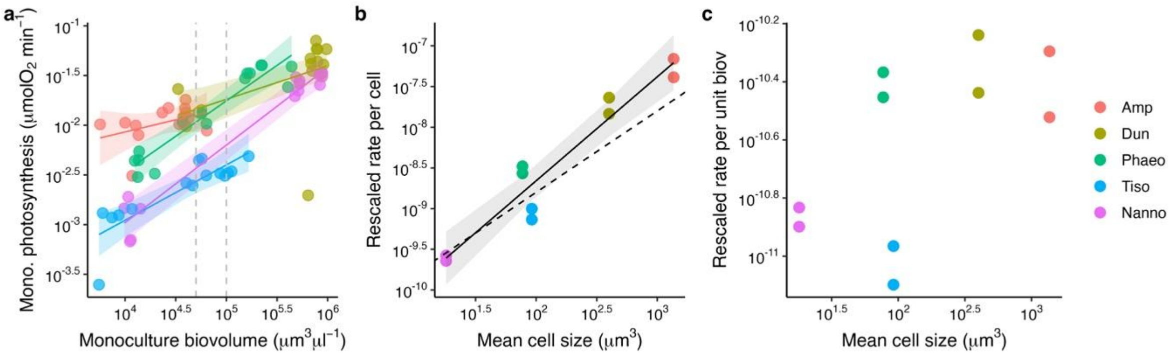


**Figure S5.** We used the relationship between photosynthesis and biovolume for each monoculture (a) to rescale the photosynthesis rate to two common biovolumes (grey lines in plot a). Comparing species at equal biovolumes shows that photosynthesis rates *per cell* increase with mean cell size with a slope not different from 1 (slope = 1.2, CI: 0.98; 1.57; b). Rates per unit biovolume, therefore, are similar among species and do not explain differences in performance between dominant (*Dunaliella*, *Nannochloropsis*) and subordinate species (c). The mean cell size shown here is the average size across the entire growth curve, averaged across all replicates of the same species.

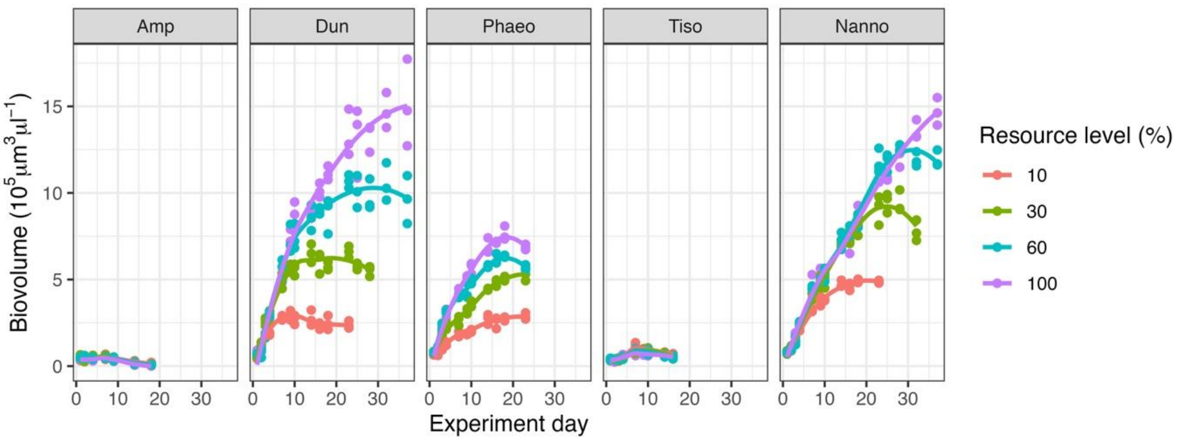


**Figure S6.** Species showed different growth responses to changes in resource levels. The dominant species (*Dunaliella* and *Nannochloropsis*) performed better across nutrient conditions than the subordinate species.

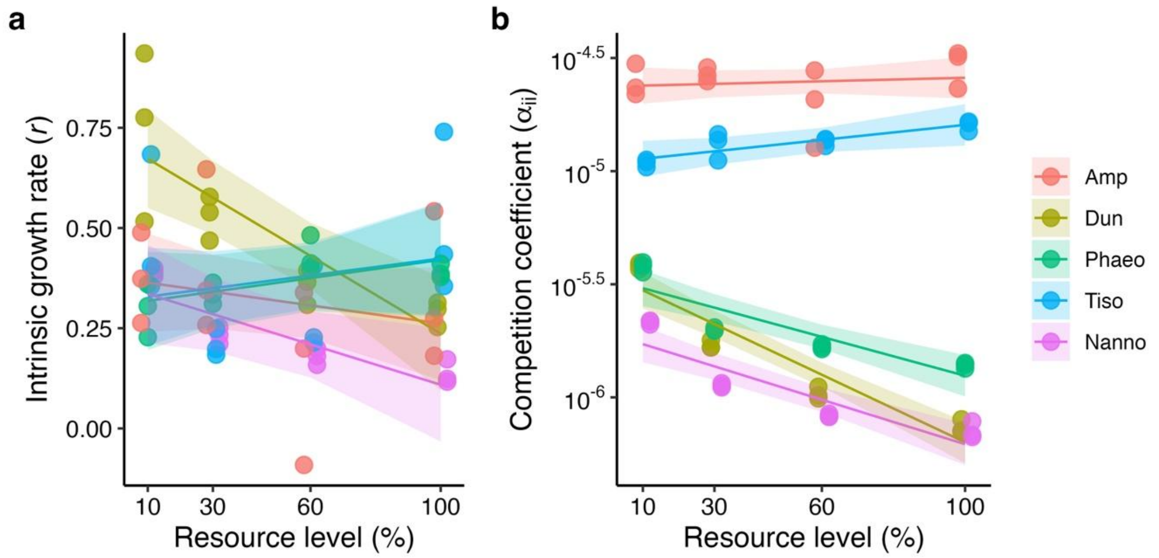


**Figure S7.** Species had different growth responses and tolerance to competition as nutrient concentration changed: dominant species (*Dunaliella*, *Nannochloropsis*) increased intrinsic growth rates ($r$) as nutrient concentration declined (a) but their sensitivity to competition ($\alpha_{ii}$) increased in parallel (b). Changes in $r$ and $\alpha_{ii}$ produce the relationships between $r$ and K showed in Figure 4f (main text).

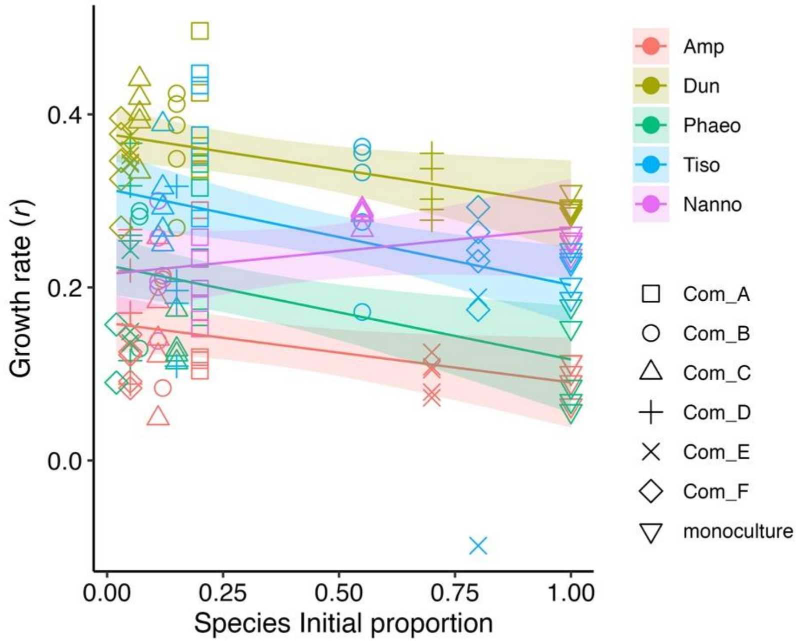


**Figure S8.** Species intrinsic growth rate (*r*) was affected by their initial proportion in the communities and monocultures. Most species showed a negative correlation between initial proportion and their intrinsic growth rate (*r*), except for *Nannochloropsis.* Abbreviations of community treatments (see also Table S1): Com_A = J1 Even; Com_B = J0.8 Tiso dominant; Com_C = J0.8 Nanno dominant; Com_D = J0.6 Dun dominant; Com_E = J0.6 Amp dominant; Com_F = J0.4 Tiso dominant.

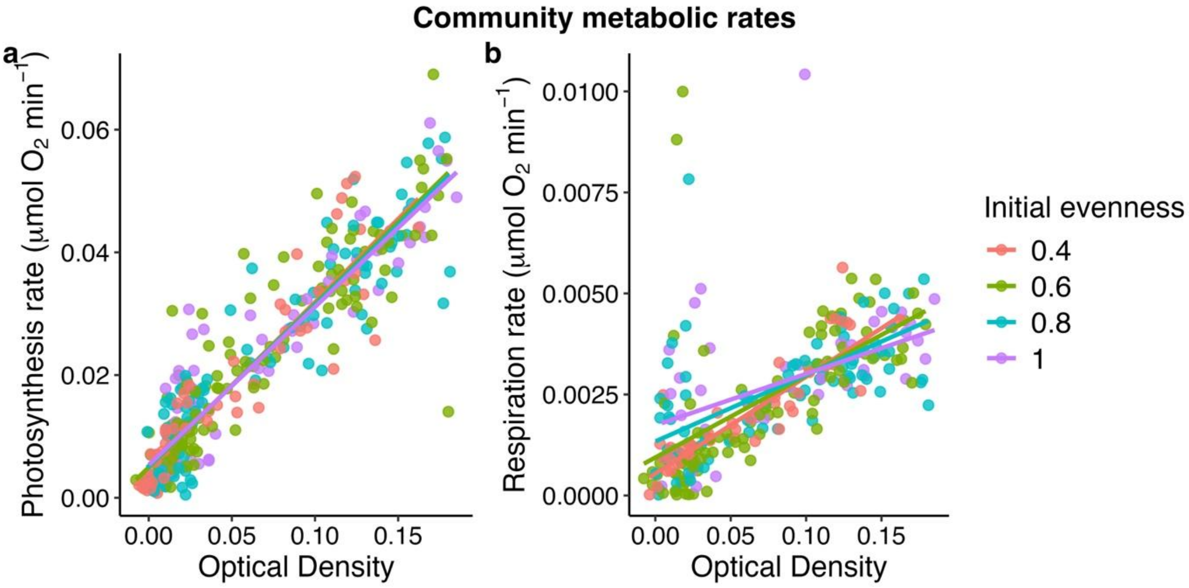


**Figure S9.** Community photosynthesis (a) increased with optical density (a proxy for biomass) independently of initial species evenness; while respiration rates (b) showed a slower increase with optical density for higher evenness values.

**Table S1:** Details of the evenness treatments and relative species proportions for each community composition ($n$ = 5).

| **Species** | **Com A:** J1 Even | **Com B:** J0.8 Tiso dominant | **Com C:** J0.8 Nanno dominant | **Com D:** J0.6 Dun dominant | **Com E:** J0.6 Amp dominant | **Com F:** J0.4 Tiso dominant |
|---|---|---|---|---|---|---|
| *Dunaliella* | 0.2 | 0.15 | 0.07 | 0.7 | 0.05 | 0.03 |
| *Tisochrysis* | 0.2 | 0.55 | 0.12 | 0.15 | 0.05 | 0.8 |
| *Phaeodactylum* | 0.2 | 0.07 | 0.15 | 0.05 | 0.05 | 0.02 |
| *Nannochloropsis* | 0.2 | 0.11 | 0.55 | 0.05 | 0.15 | 0.05 |
| *Amphidinium* | 0.2 | 0.12 | 0.11 | 0.05 | 0.7 | 0.05 |
| Shannon Diversity Index (H’) | 1.609 | 1.2968 | 1.2968 | 0.9836 | 0.9836 | 0.6615 |
| **Pielou’s Evenness Index (J)** (or Shannon Equitability Index) | **1** | **0.8057** | **0.8057** | **0.6111** | **0.6111** | **0.4110** |

| **Table S2.** Species names, strain codes and functional groups, acquired from the Roscoff Culture Collection in France. Cell volumes ($\mu m^3$) were calculated using an approximate geometric shape, either a prolate spheroid or sphere where d = diameter, h = height (length) of the cell. | | | |
|---|---|---|---|
| **Species name** | **Strain code** | **Functional group** | **Volume formula** |
| *Dunaliella tertiolecta* | RCC6 | Chlorophyta | Prolate spheroid: $V = \frac{\pi}{6} d^3$ |
| *Amphidinium carterae* | RCC88 | Dinoflagellata | |
| *Phaeodactylum tricornutum* | RCC2967 | Bacillariophyta | |
| *Nannochloropsis granulata* | RCC8 | Chlorophyta | Sphere: $V = \frac{\pi}{6} d^2 h$ |
| *Tisochrysis lutea* | RCC90 | Haptophyta | |

**Table S3.** Results of linear model estimating the effects of initial evenness (numeric predictor) and community composition (categorical, 6 levels) on the change in community evenness from the start to the end of the experiment. Significant values ($p < 0.05$) are in bold. Relates to Figure 3b.

| | **Df** | **Sum Sq** | **Mean Sq** | **F value** | **Pr(>F)** |
|---|---|---|---|---|---|
| Initial evenness | 1 | 0.00483 | 0.004828 | 0.1907 | 0.666 |
| Composition | 4 | 0.82204 | 0.205510 | 8.1150 | **0.00027** |
| Residuals | 24 | 0.60779 | 0.025325 | | |

**Table S4.** Results of linear models testing the relationship between dominance in communities (% species biovolume, $log_{10}$-transformed) and each monoculture demographic parameter (*r* or K), morphological (cell size, $\mu m^3$), or physiological trait (photosynthesis rate per unit biovolume, $\mu mol\ O_2\ min^{-1}\ \mu m^{-3}$), including an interaction with initial evenness (categorical predictor). Significant values ($p < 0.05$) are in bold.

| | Df | Sum Sq | Mean Sq | F value | Pr(>F) |
|---|---|---|---|---|---|
| Dominance ~ $r_{mono}$ × initial evenness (adj $R^2$ = 0.51) (Figure 4a) | | | | | |
| | **Df** | **Sum Sq** | **Mean Sq** | **F value** | **Pr(>F)** |
| $r_{mono}$ | 1 | 31.713 | 31.713 | 142.58 | **<0.0001** |
| Initial evenness | 3 | 0.686 | 0.229 | 1.03 | 0.38 |
| $r_{mono}$ × evenness | 3 | 0.459 | 0.153 | 0.69 | 0.56 |
| Residuals | 127 | 28.247 | 0.222 | | |
| Dominance ~ $K_{mono}$ × initial evenness (adj $R^2$ = 0.59) (Figure 4b) | | | | | |
| | **Df** | **Sum Sq** | **Mean Sq** | **F value** | **Pr(>F)** |
| $K_{mono}$ | 1 | 32.522 | 32.522 | 174.50 | **<0.0001** |
| Initial evenness | 3 | 0.182 | 0.061 | 0.33 | 0.81 |
| $K_{mono}$ × evenness | 3 | 4.732 | 0.577 | 8.46 | **<0.0001** |
| Residuals | 127 | 23.669 | 0.186 | | |
| Dominance ~ cell size × initial evenness (adj $R^2$ = 0.08) (Figure 4c) | | | | | |
| | **Df** | **Sum Sq** | **Mean Sq** | **F value** | **Pr(>F)** |
| $log_{10}$-cell size | 1 | 0.003 | 0.003 | 0.0075 | 0.93 |
| Initial evenness | 3 | 1.832 | 0.610 | 1.457 | 0.23 |
| $log_{10}$-cell size × evenness | 3 | 6.050 | 2.017 | 4.812 | **0.003** |
| Residuals | 127 | 53.220 | 0.419 | | |
| Dominance ~ photosynthesis unit volume × initial evenness (adj $R^2$ = 0.12) (Fig. 4d) | | | | | |
| | **Df** | **Sum Sq** | **Mean Sq** | **F value** | **Pr(>F)** |
| $log_{10}$-photo | 1 | 1.802 | 1.802 | 4.48 | **0.036** |
| Initial evenness | 3 | 2.128 | 0.709 | 1.76 | 0.16 |
| $log_{10}$-photo × evenness | 3 | 6.135 | 2.045 | 5.09 | **0.002** |
| Residuals | 127 | 51.04 | 0.402 | | |

**Table S5.** Results of linear model testing the effects of species biovolume ($log_{10}$-transformed) and species ID on species' photosynthetic rates ($log_{10}$-transformed). Refers to Figure 4b. Significant values ($p < 0.05$) are in bold. Refers to Figure 4e.

| Log$_{10}$ Photosynthesis rate ~ Log$_{10}$-Biov × Species ID | | | | | |
|---|---|---|---|---|---|
| | **Df** | **Sum Sq** | **Mean Sq** | **F value** | **Pr(>F)** |
| Log$_{10}$-Biovolume | 1 | 13.1969 | 13.1969 | 269.8492 | **< 0.0001** |
| Species ID | 4 | 5.7579 | 1.4395 | 29.4342 | **< 0.0001** |
| Log$_{10}$-Biov × Sp ID | 4 | 0.8536 | 0.2134 | 4.3637 | **0.003497** |
| Residuals | 64 | 3.1299 | 0.0489 | | |

| Post-hoc results: species slopes | | | | | |
|---|---|---|---|---|---|
| Species ID | Slope | SE | df | Lower CL | Upper CL |
| *Amphidinium* | 0.245 | 0.197 | 64 | -0.150 | 0.639 |
| *Dunaliella* | 0.338 | 0.105 | 64 | 0.127 | 0.548 |
| *Nannochloropsis* | 0.780 | 0.068 | 64 | 0.644 | 0.917 |
| *Phaeodactylum* | 0.715 | 0.103 | 64 | 0.507 | 0.919 |
| *Tisochrysis* | 0.557 | 0.115 | 64 | 0.328 | 0.786 |

| Post-hoc results: species intercepts at biov = $10^{4.9}$ (EMM = estimated marginal mean) | | | | | |
|---|---|---|---|---|---|
| Species ID | EMM | SE | df | Lower CL | Upper CL |
| *Amphidinium* | -1.85 | 0.116 | 64 | -2.08 | -1.62 |
| *Dunaliella* | -1.77 | 0.089 | 64 | -1.95 | -1.60 |
| *Nannochloropsis* | -2.28 | 0.062 | 64 | -2.40 | -2.16 |
| *Phaeodactylum* | -1.83 | 0.058 | 64 | -1.94 | -1.71 |
| *Tisochrysis* | -2.46 | 0.07 | 64 | -2.60 | -2.32 |

**Table S6.** Results of linear model testing the effect of monocultures biomass (optical density, OD) on population photosynthesis rates ($log_{10}$-transformed). Significant values ($p < 0.05$) are in bold. Refers to Figure S4.

| Log$_{10}$ Photosynthesis rate ~ log$_{10}$ optical density × species | | | | | |
|---|---|---|---|---|---|
| | **Df** | **Sum Sq** | **Mean Sq** | **F value** | **Pr(>F)** |
| Log$_{10}$-Optical Density | 1 | 48.883 | 48.88 | 582.50 | **<0.0001** |
| Species ID | 4 | 7.595 | 1.899 | 22.625 | **<0.0001** |
| Log$_{10}$-OD × Species ID | 4 | 1.944 | 0.486 | 5.7907 | **0.000167** |
| Residuals | 303 | 25.427 | 0.084 | | |
| Post-hoc results: species photosynthesis slopes | | | | | |
| Species ID | Slope | SE | df | Lower CL | Upper CL |
| *Amphidinium* | 0.748 | 0.4032 | 303 | -0.0453 | 1.542 |
| *Dunaliella* | 0.587 | 0.0636 | 303 | 0.4622 | 0.713 |
| *Nannochloropsis* | 0.781 | 0.0759 | 303 | 0.6319 | 0.931 |
| *Phaeodactylum* | 0.696 | 0.0479 | 303 | 0.6020 | 0.790 |
| *Tisochrysis* | 0.343 | 0.0698 | 303 | 0.2054 | 0.480 |
| Post-hoc results: species intercepts at log$_{10}$-OD = -1.61 (EMM = estimated marginal mean) | | | | | |
| Species ID | EMM | SE | df | Lower CL | Upper CL |
| *Amphidinium* | -2.07 | 0.0421 | 303 | -2.16 | -1.99 |
| *Dunaliella* | -1.89 | 0.0398 | 303 | -1.96 | -1.81 |
| *Nannochloropsis* | -2.24 | 0.0374 | 303 | -2.31 | -2.17 |
| *Phaeodactylum* | -2.21 | 0.0365 | 303 | -2.28 | -2.14 |
| *Tisochrysis* | -2.46 | 0.0426 | 303 | -2.54 | -2.37 |

**Table S7.** Results of linear model estimating the effects of resource level (nutrient percentage as a numeric predictor) and Species ID (categorical) on intrinsic growth rates (*r*). Significant values ($p < 0.05$) are in bold. Refers to Figure S7a.

| Max. growth rate (*r*) ~ Resource level × Species ID | | | | | |
|---|---|---|---|---|---|
| | **Df** | **Sum Sq** | **Mean Sq** | **F value** | **Pr(>F)** |
| Resource level | 1 | 0.10951 | 0.109515 | 5.8630 | **0.019133** |
| Species ID | 4 | 0.37716 | 0.094290 | 5.0479 | **0.001700** |
| Resource level × Species ID | 4 | 0.35316 | 0.088289 | 4.7267 | **0.002589** |
| Residuals | 50 | 0.93395 | 0.018679 | | |

| Post-hoc results: species slopes | | | | | |
|---|---|---|---|---|---|
| Species | Slope | SE | df | Lower CL | Upper CL |
| *Amphidinium* | -0.00113 | 0.00116 | 50 | -0.00347 | 0.001204 |
| *Dunaliella* | -0.00485 | 0.00116 | 50 | -0.00718 | -0.002511 |
| *Phaeodactylum* | 0.00114 | 0.00116 | 50 | -0.00120 | 0.003476 |
| *Tisochrysis* | 0.00106 | 0.00116 | 50 | -0.00128 | 0.003395 |
| *Nannochloropsis* | -0.00252 | 0.00116 | 50 | -0.00485 | -0.000179 |

**Table S8.** Results of linear model estimating the effects of resource level (nutrient percentage as a numeric predictor) and Species ID (categorical) on species density-dependence ($\alpha_{ii}$, $\log_{10}$-transformed). Significant values ($p < 0.05$) are in bold. Refers to Figure S7b.

Density-dependence ($\alpha_{ii}$) ~ Resource level × Species ID

| | **Df** | **Sum Sq** | **Mean Sq** | **F value** | **Pr(>F)** |
|---|---|---|---|---|---|
| Resource level | 1 | 0.59 | 0.59 | 75.045 | **< 0.0001** |
| Species ID | 4 | 17.76 | 4.44 | 564.13 | **< 0.0001** |
| Resource level × Species ID | 4 | 0.81 | 0.20 | 25.637 | **< 0.0001** |
| Residuals | 50 | 0.39 | 0.001 | | |

Post-hoc results: species slopes

| Species | Slope | SE | df | Lower CL | Upper CL |
|---|---|---|---|---|---|
| *Amphidinium* | 0.00038 | 0.0008 | 50 | -0.0011 | 0.0019 |
| *Dunaliella* | -0.00747 | 0.0008 | 50 | -0.009 | -0.006 |
| *Phaeodactylum* | -0.00430 | 0.0008 | 50 | -0.0058 | -0.0028 |
| *Tisochrysis* | 0.001667 | 0.0008 | 50 | 0.00015 | 0.00318 |
| *Nannochloropsis* | -0.00492 | 0.0008 | 50 | -0.00643 | -0.0034 |

**Table S9.** Results of linear model estimating the effects of intrinsic growth rate (*r*) (as a numeric predictor) and Species ID (categorical) on species carrying capacity (K) across different resource levels. Significant values ($p < 0.05$) are in bold. Refers to Figure 4f.

| Carrying capacity (K) ~ *r* × Species ID | | | | | |
|---|---|---|---|---|---|
| | **Df** | **Sum Sq** | **Mean Sq** | **F value** | **Pr(>F)** |
| Intrinsic growth rate (*r*) | 1 | 9.98e+11 | 9.98e+11 | 59.487 | **< 0.0001** |
| Species ID | 4 | 8.59e+12 | 2.15e+12 | 128.029 | **< 0.0001** |
| *r* × Species ID | 4 | 2.12e+12 | 5.30e+11 | 31.575 | **< 0.0001** |
| Residuals | 50 | 8.38e+11 | 1.68e+10 | | |

| Post-hoc results: species slopes | | | | | |
|---|---|---|---|---|---|
| Species | Slope | SE | df | Lower CL | Upper CL |
| *Amphidinium* | -34545 | 204775 | 50 | -445848 | 376757 |
| *Dunaliella* | -1771450 | 188556 | 50 | -2150176 | -1392725 |
| *Phaeodactylum* | 1814085 | 604744 | 50 | 599420 | 3028750 |
| *Tisochrysis* | -1345 | 214125 | 50 | -431428 | 428737 |
| *Nannochloropsis* | -3606533 | 384295 | 50 | -4378413 | -2834653 |

**Table S10.** Results of linear model estimating the effects of species initial proportion (numeric predictor) and Species ID (categorical) on species intrinsic growth rate (*r*) in the communities and monoculture. Significant values ($p < 0.05$) are in bold. Refers to Fig. S8.

| Intrinsic growth rate (*r*) ~ Species Initial Proportion × Species ID | | | | | |
|---|---|---|---|---|---|
| | **Df** | **Sum Sq** | **Mean Sq** | **F value** | **Pr(>F)** |
| Initial Proportion | 1 | 0.07277 | 0.072767 | 14.4884 | **0.0002119** |
| Species ID | 4 | 0.87293 | 0.218233 | 43.4519 | **<0.0001** |
| Initial Prop. × Species ID | 4 | 0.04911 | 0.012277 | 2.4445 | **0.0494787** |
| Residuals | 137 | 0.68807 | 0.005022 | | |

| Post-hoc results: species slopes | | | | | |
|---|---|---|---|---|---|
| Species | Slope | SE | df | Lower CL | Upper CL |
| *Amphidinium* | -0.0688 | 0.0347 | 137 | -0.1374 | -0.000214 |
| *Dunaliella* | -0.0822 | 0.0340 | 137 | -0.1494 | -0.015007 |
| *Phaeodactylum* | -0.1089 | 0.0386 | 137 | -0.1852 | -0.032700 |
| *Tisochrysis* | -0.1111 | 0.0370 | 137 | -0.1842 | -0.037941 |
| *Nannochloropsis* | 0.0532 | 0.0453 | 137 | -0.0363 | 0.142794 |

**Table S11.** Results of linear model testing the effects of community biomass (optical density, OD) and initial evenness (both numeric predictors) on communities' metabolic rates. Significant values ($p < 0.05$) are in bold.

| | | | | | |
|---|---|---|---|---|---|
| Photosynthesis rate ~ optical density × initial evenness (Figure S9a) | | | | | |
| | **Df** | **Sum Sq** | **Mean Sq** | **F value** | **Pr(>F)** |
| OD | 1 | 0.078068 | 0.078068 | 1999.25 | **<0.0001** |
| Initial evenness | 1 | 0.000001 | 0.000001 | 0.03 | 0.8648 |
| OD × evenness | 1 | 0.000021 | 0.000021 | 0.53 | 0.4691 |
| Residuals | 367 | 0.014331 | 0.000039 | | |
| Respiration rate ~ optical density × initial evenness (Figure S9b) | | | | | |
| | **Df** | **Sum Sq** | **Mean Sq** | **F value** | **Pr(>F)** |
| OD | 1 | 0.00026 | 2.6457e-04 | 161.83 | **<0.0001** |
| Initial evenness | 1 | 0.0000031 | 3.1210e-06 | 1.91 | 0.168 |
| OD × evenness | 1 | 0.0000096 | 9.6010e-06 | 5.87 | **0.016** |
| Residuals | 258 | 0.0004218 | 1.6350e-06 | | |
